# Chromatin Remodeling and Epigenetic Modifications in Planar Cell Migration

**DOI:** 10.1101/2023.12.21.572681

**Authors:** Jack Forman, Briar Hine, Samantha Kaonis, Soham Ghosh

## Abstract

The cell nucleus plays a critical role in cell migration by its deformability under forces, by acting as a piston that activates mechanosensitive channels, and also by serving as a ruler tailoring cell response to spatial constraints. Cell nuclear mechanics determine the mechanobiology of static and migrating cells. Here, we report that intranuclear chromatin architecture plays a previously unknown role during cell migration on planar substrates. By inhibiting histone deacetylation, which compacts chromatin, and a methyltransferase, which affects chromatin remodeling and chromatin compaction, we showed that cell migration speed and persistence drastically slowed wound closure efficiency in an in vitro scratch wound assay. Chromatin remodeling was visualized and quantified during cell migration, which may be intertwined with nuclear mechanics and shape. Inhibition disrupts remodeling and other unknown roles in chromatin, thus affecting the speed and persistence of cell migration. Interestingly, cytoskeletal stress fiber formation and cell shape were not visibly affected by chromatin modifications, suggesting an exclusive nuclear mechanobiological role in cell migration. These findings provide new insights into how aging and other degenerative conditions affect the plasticity of chromatin architecture and hence its effect on cell migration efficiency.

## Introduction

Cell migration is a fundamental process in normal physiology and pathology that affects wound healing, developmental biology, cancer cell metastasis, and cell infiltration inside scaffolds during tissue engineering. Cells migrate in response to external chemicals or physical stimuli. During this process, the actin cytoskeleton is generally referred to as the engine of motility because of its elaborate network, which drives the first step of cell movement through protrusion, using forces generated from asymmetric polymerization. Recently, the nucleus has emerged as a critical mediator of cell migration (1). Nuclear movement and positioning, along with deformability, nuclear envelope content, and nucleus-cytoskeleton connectivity, have been shown to be determining factors during cell migration (2). Experiments with enucleated cells called cytoblasts revealed that in 1D or 2D migration, nuclei are dispensable for migration, but critical for proper cell mechanical responses (3). Cell migration in confined environments has received particular attention in recent years, and it has been shown that DNA damage occurs via large deformations in constricted spaces, which are overcome by cancer cells because of their adaptability (4). More recently, studies have shown that the nucleus serves as a ruler tailoring the cell response to spatial constraints (5) and acts as a piston that activates mechanosensitive channels (6, 7). All these studies clearly indicate that the roles of the nucleus and its mechanics in cell migration are yet to be understood.

In previous studies, bulk nuclear mechanics and shape were considered while understanding the nuclear role of cell migration; however, the role of chromatin-level mechanics and its dynamic nature in cell migration has not been investigated so far. Newly discovered studies point towards the emerging roles of intranuclear chromatin level heterogeneity in cell functions (8). Accordingly, some investigations have shown that intranuclear mechanics can be spatially variable (9–11), which has implications for how mechanical force is experienced by the cell nucleus. Because cells experience heterogeneous levels of cytoskeletal forces at different cell locations, it is possible that different locations inside the nucleus experience different amounts of force and chromatin movement to coordinate local nuclear mechanics with local cell mechanics, as shown in chondrocyte deformation during osmotic loading (12).

The intranuclear space of the cell is dynamic, and chromatin remodeling has emerged as a means for the nucleus to differentially express genes over time. Such remodeling occurs through epigenetic mechanisms, including ATP-dependent chromatin remodeling and histone modifications that locally condense or decondense the chromatin (13). The detailed mechanism of chromatin remodeling and its functional significance has only recently emerged. Histone modifications that occur at the nucleosome level drive chromatin compaction by wrapping or unwrapping DNA around nucleosomes. Histone deacetylases (HDACs) mediate a subtype of such modifications. Inhibition of HDAC hinders chromatin compaction and keeps the chromatin open, thus rendering the nucleus softer (14). If chromatin remodeling at the nucleosome level by wrapping or unwrapping is critical for cell motility, migration would be affected by HDAC inhibition. Another type of chromatin remodeling occurs at the multi nucleosome level, where ATP-dependent chromatin remodelers are critical (15). The precise mechanism of chromatin remodeling that occurs through nucleosome sliding, engagement, and ejection is still being discovered; however, several classes of chromatin remodeling complexes have been discovered, including SWI/SNF (16). The core protein of SWI/SNF is ARID1A, which maintains a delicate balance with EZH2. ARID1A, as a part of SWI/SNF, determines chromatin remodeling; hence, inhibiting EZH2 can potentially intervene with the chromatin remodeling process. Therefore, pharmacological intervention with HDAC and EZH2 provides an opportunity to investigate the role of chromatin remodeling in cell functions, including cell migration. Altered chromatin mechanics is also clinically relevant, as studies have found that with aging, nuclear mechanics changes due to the altered state of epigenetic modifications in the cell nucleus (17).

In this study, we hypothesized that chromatin remodeling during cell migration and interfering with chromatin remodeling can affect cell migration speed and persistence. To test this hypothesis, we quantified cell migration using a scratch wound assay in NIH 3T3 cells, along with high-resolution measurement of chromatin remodeling inside the cell nucleus. Chromatin remodeling is inhibited by two different pharmacological interventions: (1) inhibition of HDAC with Trichostatin A (TSA) and (2) inhibition of methyltransferase EZH2, which regulates ARID1A activity and chromatin compaction with GSK126. Instead of disrupting nuclear mechanics completely by a large dosage of pharmacological treatments or by nucleo-cytoskeletal decoupling, we applied a small amount of drug that did not visibly affect the cell phenotype or viability. The effects of these treatments on cell migration, chromatin remodeling, and nuclear shape were quantified. The results are discussed to provide novel insights into the role of chromatin remodeling in planar cell migration, with further possible implications for 3D migration.

### Materials and Methods

#### Cell culture and scratch wound assay

The murine fibroblast cell line NIH 3T3 (a gift from Corey Neu Lab at the University of Colorado Boulder) was used for all experiments. Briefly, the cells were maintained in the culture medium made of DMEM (ATCC 30-2002), penicillin/streptomycin (Gibco, 15140-122) and Fetal Bovine Serum (EqualFETAL, Atlas Biologicals). Trypsinization for cell passaging and other purposes was performed using TrypLE (Thermo Fisher, 12605010). The scratch wound assays used 8 well µ-Slide plate (ibidi, 80821). To functionalize these plates for cell attachment, the wells were plasma-treated for a few seconds, followed by the application of bovine Type 1 collagen solution (Gibco, A10664) for at least 1 hour. Subsequently, the cells were seeded into wells and maintained in medium. After an incubation period of 48 h, the cell layer confluency was observed. A scratch was applied to the cell monolayer using a 200 μL pipette tip guided by a straightedge. Next, the wells were quickly rinsed with DPBS to remove any debris and were provided with fresh cell culture medium.

#### Pharmacological treatment of cells

Drugs were used to induce chromatin modifications in the cells. Drugs were applied 14 h before scratching was induced. While the medium was changed after the scratch was made, the change was made in the control (no drug) group too. The drug-treated groups received fresh medium containing drugs after scratching. The following two drug treatment groups were used: GSK126 (Sigma) at 20 μM and TSA (Sigma) at 100 ng/ml. GSK126 is a methyltransferase EZH2 inhibitor that affects the balance of ARID1A in SWI/SNF, thereby affecting chromatin remodeling and lowering facultative chromatin compaction. TSA is a histone deacetylase (HDAC) inhibitor that lowers chromatin condensation.

#### Imaging of live cell migration and the fixed cells post migration

All imaging was done using a Zeiss LSM 980 inverted confocal microscope. For live-cell imaging, brightfield and epifluorescence modes were used. Nuclei in live cells were stained with Nucblue (Thermo Fisher, R37605) 30 min before scratching was applied. For live imaging, 8 well slides were placed inside the microscope with a sufficient amount of culture medium so that the medium did not dry out. Physiologically relevant temperature (37° °C), CO_2_ concentration (5%), and humidity levels were maintained in the microscope using an incubation chamber. The tile mode of the microscope was used for time-lapse imaging of a larger field of view. Staining of fixed cells was performed using DAPI (Thermo Fisher, D1306) and phalloidin (Thermo Fisher, R37110) after fixing with 4% paraformaldehyde, permeabilization with Triton X, and blocking of non-specific binding sites with bovine serum albumin. High-resolution and high-magnification images of individual nuclei in the migrating cells were obtained using a 63× oil objective in separate experiments.

#### Image analysis for the quantification of cell migration parameters

All image analyses were performed using the open-source software Fiji. First, the percentage of wound closure at the given time points was calculated. This was done by outlining a region of interest (ROI) of the initial scratch in the brightfield images and calculating an area measurement serving as A_0_. At a later time point, the total area (A_t_) of cells in the initial ROI was calculated. The percentage of wound closure was then calculated as 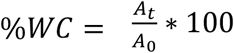 . To quantify the cell-specific migration parameters, the Fiji plugin TrackMate was applied. The resulting migration tracks allowed for the calculation of several parameters, including the total distance travelled, mean speed, linearity of forward progression, and instantaneous angle of the cell trajectory at a specific time point.

#### Chromatin remodeling quantification from high resolution images

High-resolution z-stack images were projected onto a single plane for 2D analysis of the images. An in-house image-based registration technique was used to correlate nuclear images at any pair of time points. Briefly, images were corrected for translation and rotation. Then, the texture of the images was used to correlate the template and target images that provided the displacement map in the nucleus. This technique is limited only by the image resolution. We defined the displacement map as a chromatin remodeling map for this specific context. This technique has been thoroughly validated in previous studies (18).

#### Statistics

ANOVA followed by post-hoc tests was used to quantify differences between the groups. Error bars in bar graphs represent the standard deviation. The number of technical replicates, number of cells (when applicable), and p-values are reported in each individual figure.

## Results

### Chromatin modifications impair the wound closure efficiency

A scratch wound assay was developed and validated as reported in numerous previous studies. The wounds in the control groups were 80% closed by 11 h and completely closed by 24 h (Figure 1). Further validation of this assay was performed using Y27632, a ROCK inhibitor that impairs actomyosin contractility, thereby slowing cell migration (data not shown). The control group shows that after 24 h, the wound is completely closed. Two chromatin modification strategies were used for the application of TSA and GSK126. TSA impairs chromatin condensation in the nucleus via HDAC inhibition which condenses the chromatin. TSA application (100 ng/mL) showed that after 24 h, only approximately 30% of the wound area was closed by migrating cells. GSK126 impairs chromatin remodeling by inhibiting EZH2, which directly affects chromatin remodeling by disrupting the dynamics of ARID1A, the key component of the chromatin remodeler SWI/SNF, or by affecting H3K27 methylation, which condenses chromatin. GSK126 application showed that after 24 h, only approximately 25% of the wound area was closed by migrating cells.

**Figure 1.**
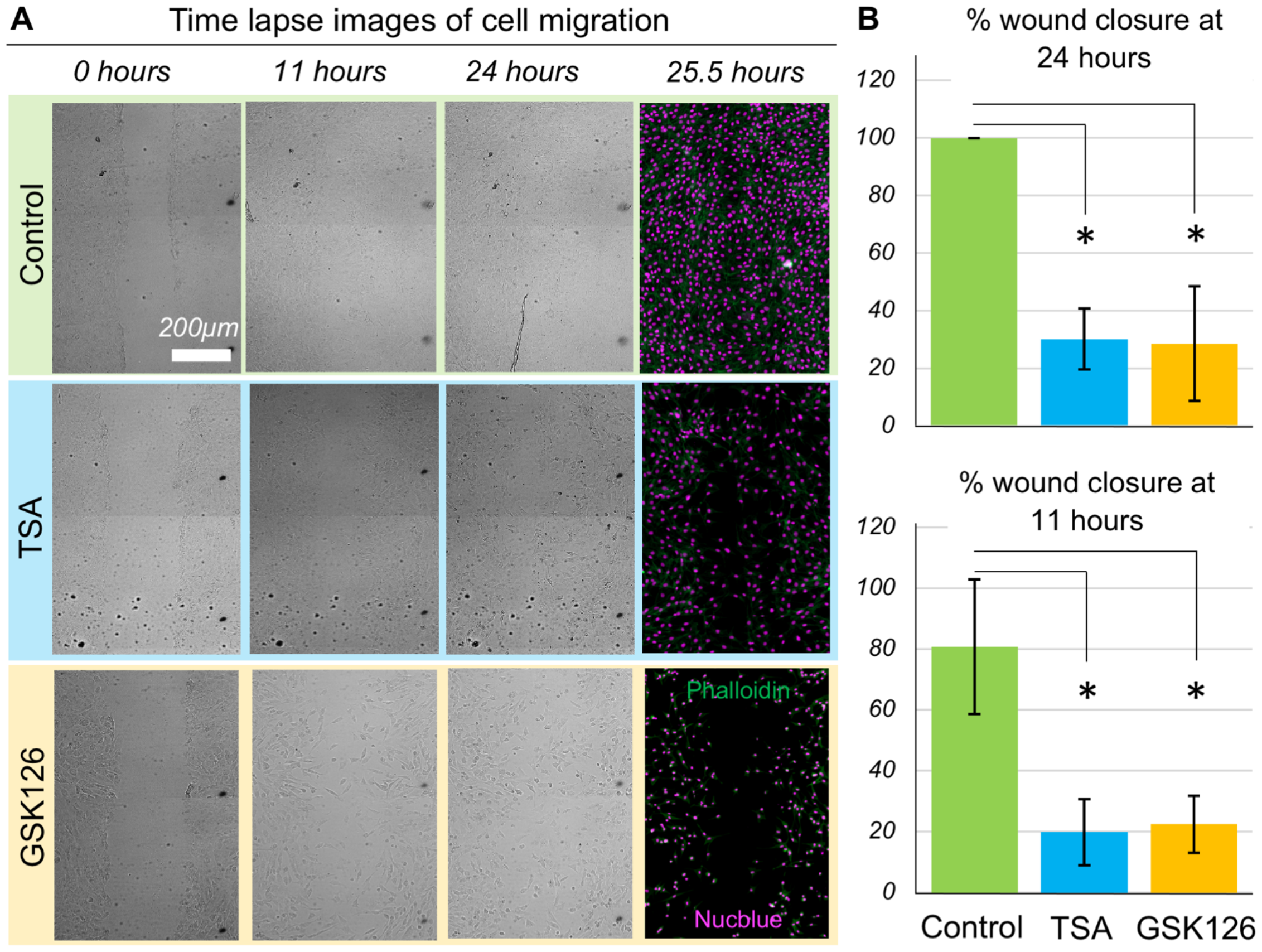
Wound closure is affected by the chromatin modification. **(A)** Scratch wound assay with NIH 3T3 cells show that they migrate to close the scratch completely by 24 hours which is impaired by the modification of the chromatin architecture and mechanics. Bright field images of live cells shown at 0 hour (right after the scratch), at 11 hours and at 24 hours timepoints. The 25.5 hour - timepoint image represents the same field of view after fixing and staining for the DNA and the F-actin. **(B)** Initial and final areas of the wound quantify the percent wound closure over time. Chromatin modifications drastically impair the wound closure efficiency. Data based upon >10 samples per group. *p < 0.01.

### Chromatin modifications disrupt the speed of individual cell migration

To understand how the individual cell migration trajectory impairs wound closure efficiency, the speed of individual cells was calculated. While the control group cells showed that over time, they persistently moved, often in groups (Figure 2A), in the GSK126 and TSA groups, only a few cells started moving from the scratch boundary. The average speed over the complete period of migration revealed that both GSK126 and TSA groups displayed almost 50% speed reduction (Figure 2B) and, hence, almost 50% total displacement (Figure 2C) over 10.5 hours compared to the control group. It was also observed that cells initially came out of the cell cluster relatively quickly to scratch, after which their speed decreased in both the TSA and GSK126 groups.

**Figure 2.**
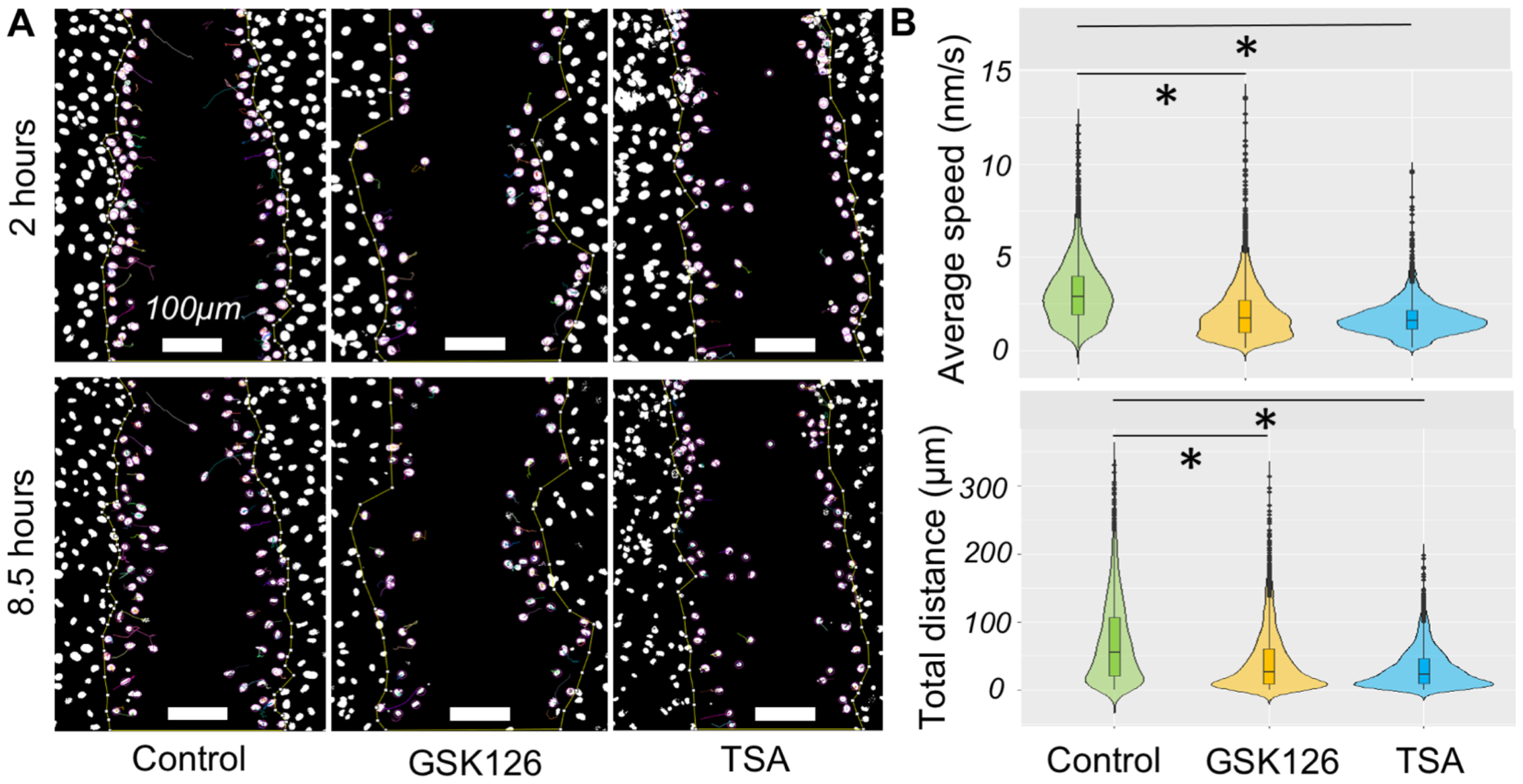
Characteristics of the kinematics of cell migration upon chromatin modification. **(A)** Tracked paths of individual nuclei at two timepoints (2 hours and 8.5 hours) are shown. For the same field of view GSK126 ad TSA groups show less cell movement. **(B)** Overlayed boxplot and violin plot of average speed and total distance traveled. Both average speed and total distance traveled (over 10.5 hours) show that chromatin modifications significantly lower the migration velocity. *p< 0.01 based upon >1800 tracked nuclei per group from at least total 6 samples per group.

### Persistence of the cell migration is reduced by the chromatin modifications

Next, the linearity of the cell path and angular orientation of cells were quantified during their migration to understand how persistently they followed the chemical gradient to close the wound. If the cells perfectly follow a straight line along the direction of the chemical gradient, the linearity is 1. If they change direction continuously, the value is closer to zero (Figure 3A). The linearity of cell migration decreased slightly, but by a statistically significant amount, in both GSK126 and TSA groups compared to the control group (Figure 3B). Ideally, if they all move perpendicular to the cell monolayer boundary and follow the chemical gradient during the entire migration trajectory, the angle should be 0º (Figure 3A). There would be some variability in this statement, although scratch creation is not a perfectly straight line in the field of view. It was found that both at individual time points and overall, at all time points, the control group had more cells near lower angles closer to 0º, and fewer cells at larger angles between 90º and 180º. A cell that moved completely in the reverse direction was designated to be 180º. Both the GSK126 and TSA groups showed that more cells were clustered in the bins at higher angles, suggesting a lack of persistent linear movement of cells along the direction of the chemical gradient (Figure 3C). The lack of persistent linear movement is further demonstrated when sampling individual time points such as 7 h and 10.5h. This is because on average the drug treated cells move along the concentration gradient and close the wound, as seen in Figure 1. However, when observing shorter time segments the cells’ ability to follow an optimum trajectory is inhibited, seen by the flatter distributions of the treated cells at single time points.

**Figure 3.**
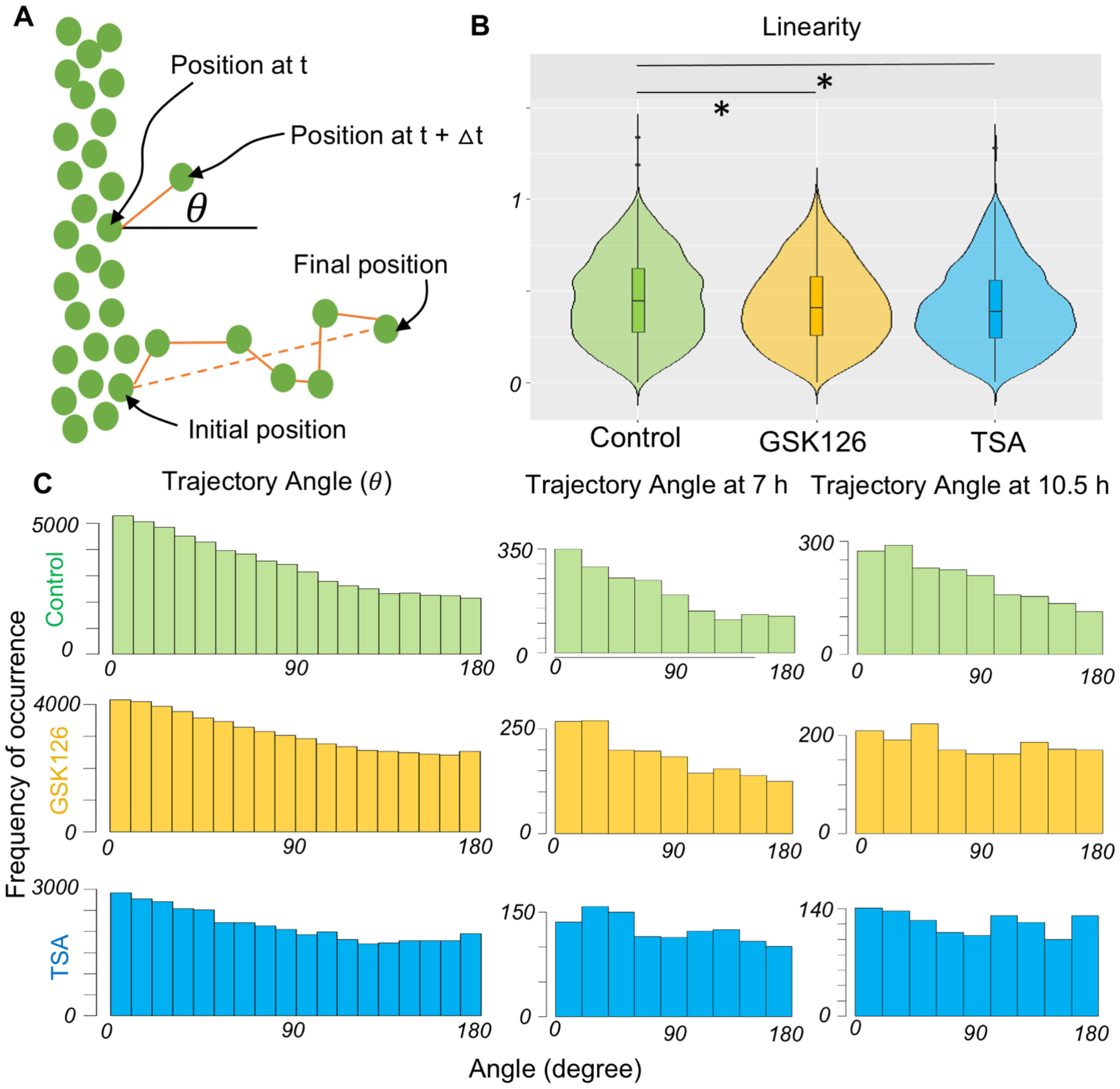
Characteristics of the dynamics of cell migration upon chromatin modification. **(A)** The persistence of cell migration directionality can be measured by the linearity and angle of cell migration direction. The schematic explains the angle of the direction (*θ*) of a migrating cell compared to the horizontal axis which signifies the ideal direction (0º) of cell migration in the scratch. Linearity is defined by the ratio of the straight dotted line between the initial and final position of the cell over the actual path traversed by the cell. **(B)** Linearity is slightly decreased by the chromatin modification signifying a cell does not follow the ideal migration path. An ideal linearity is 1. **(C)** The frequency distribution of cell count for a specific angle range for many cells in a sample for all timepoints (10.5 hours) shown in the left panel. The ideal angle of migration is 0º whereas 180º means the cell is migrating opposite to the direction of the chemical gradient. More cell counts are clustered around the lower angle in the control group. Chromatin modifying drugs make the cell migration direction more random in many non-ideal directions as shown by the histogram at all timepoints and specific timepoints (7 h and 10.5 h). *p< 0.01 based upon >1800 tracked nuclei per group from at least total 6 samples per group.

### Gross cell phenotype is not affected by the chromatin modifier drugs

Cell migration is traditionally determined by the actin cytoskeleton and actomyosin contraction. The continuous formation of the leading and trailing edges is determined by cytoskeletal F-actin turnover. Therefore, F-actin structure was investigated for the application of chromatin-modifying drugs. Surprisingly, the actin structure was mostly intact (Figure 4) upon application of the drugs. Even stress fibers were clearly visible in the migrating and non-migrating cells. It is possible that the overall nuclear mechanics might be affected by drugs (not measured in this study), which suggests an exclusive nuclear mechanical role in cell migration. However, no visible change in intranuclear chromatin architecture was observed upon application of chromatin-modifying drugs. The formation of euchromatin and heterochromatin foci was not drastically different between the groups, which led to further investigate the nuclear shape and chromatin architecture in migrating cells.

**Figure 4.**
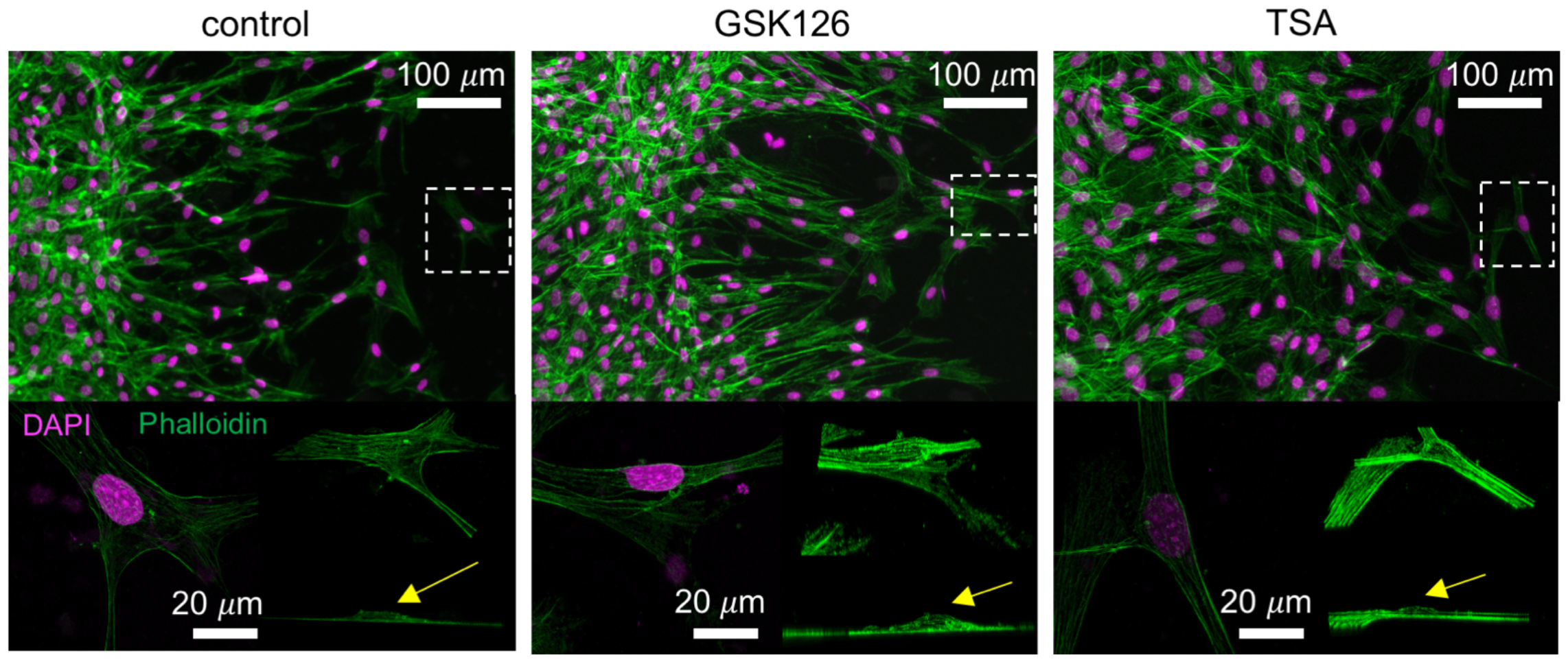
Characteristics of the stress fiber formation in migrating cells upon chromatin modification. The staining of NIH 3T3 cells reveal that F-actin or stress fiber formation is not visibly disrupted by the chromatin modifying drugs. Cells are migrating from left to right. Bottom panel shows a higher magnification images of a selected cell in the field of view (in white dotted box) along with its 3D isometric view and side view. The nuclear dome (yellow arrow) and the actin stress fiber formation around the nucleus is visible.

### Chromatin remodels during the cell migration which is impaired by the chromatin modifying drug GSK126

Intranuclear architecture is not static, but rather dynamic even on a smaller timescale, such as minutes. Next, we investigated whether the nucleus only moves with the cell or whether intranuclear chromatin also remodels during the process. From the time-lapse data, it is evident that the nuclei change their shape during the migration process (Figure 5, 6). For the control group, we found that nuclei indeed changed shape, as shown by the changing area, but the nuclear integrity, as demonstrated by the shape, remained intact (Figure 5A, B). It is possible that cytoskeletal remodeling deforms the nucleus during cell migration, but the intranuclear chromatin remodeling map revealed an interesting role of dynamic chromatin in this process (Figure 5C, 6C). We found chromatin remodeling inside the nucleus as shown by the heterogeneous color map in the nucleus. Some bulk deformation of the nucleus is also captured in the color map, which is probably caused by the force created by cytoskeletal reorganization, as shown by the red periphery in the nuclear envelope. However, this displacement also contributed to internal chromatin remodeling, which was more evident in the interior of the nucleus (Figure 5C, 6C). Upon the application of GSK126, nuclear shape and integrity were disrupted (Figure 6A, B), although chromatin remodeling was still evident inside the nucleus. The quiver plots show the direction of chromatin remodeling at a particular time instant and show a high degree of heterogeneity inside the cell nucleus, which further confirms that internal chromatin remodeling happens, independent of bulk nuclear deformation (Figure 5D, 6D).

**Figure 5.**
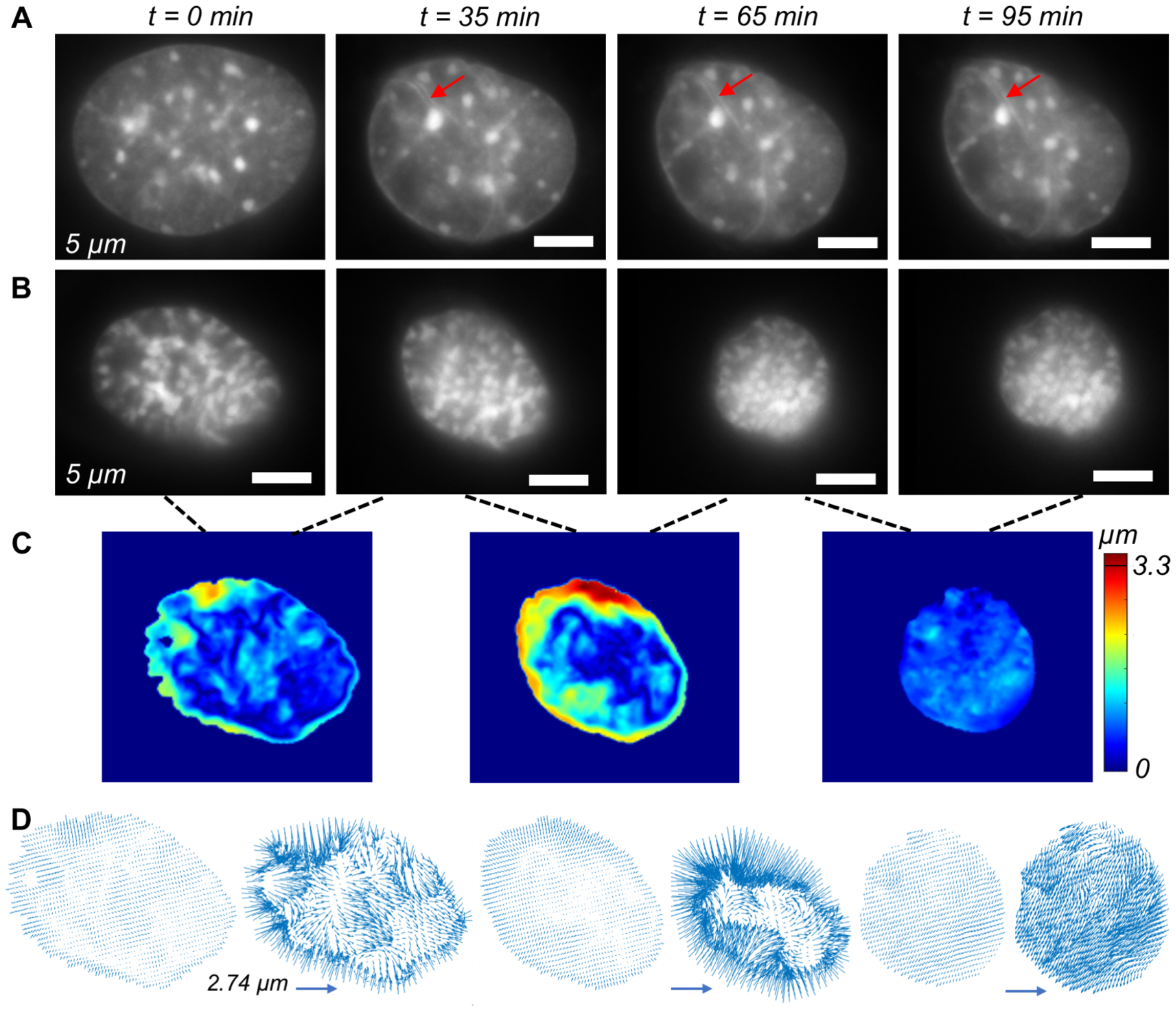
Single cell level chromatin tracking of individual Nucblue stained nuclei in the control group of NIH 3T3 cells. In all cases, the yellow arrow signifies the approximate direction of the cell migration at that timepoint. **(A)** Snapshots of a single nucleus shows that it undergoes intranuclear shape change as shown by the change in the nuclear shape and area. Wrinkling of the nucleus is visible in several regions (red arrow). **(B)** Another example of a nucleus undergoing more extreme shape and area change. In both cases, nuclear ruffling (irregularities) is not visible in the xy plane which is the plane shown the images. **(C)** Rigid body motion corrected absolute displacement map of the chromatin shows the chromatin remodeling at different pairs of timesteps i.e., 0 to 35 min, 35 min to 65 min, and 65 min to 95 min. **(D)** Vector map of the displacement field shows significant heterogeneity in the intranuclear space showing how the chromatin flows to place the nucleus inside the nucleus during the cell migration. The vector map at the right corresponding to the scale bar. The vector map in left shows scaled plots to show the exact location of the vectors, but do not represent the actual displacement magnitude.

**Figure 6.**
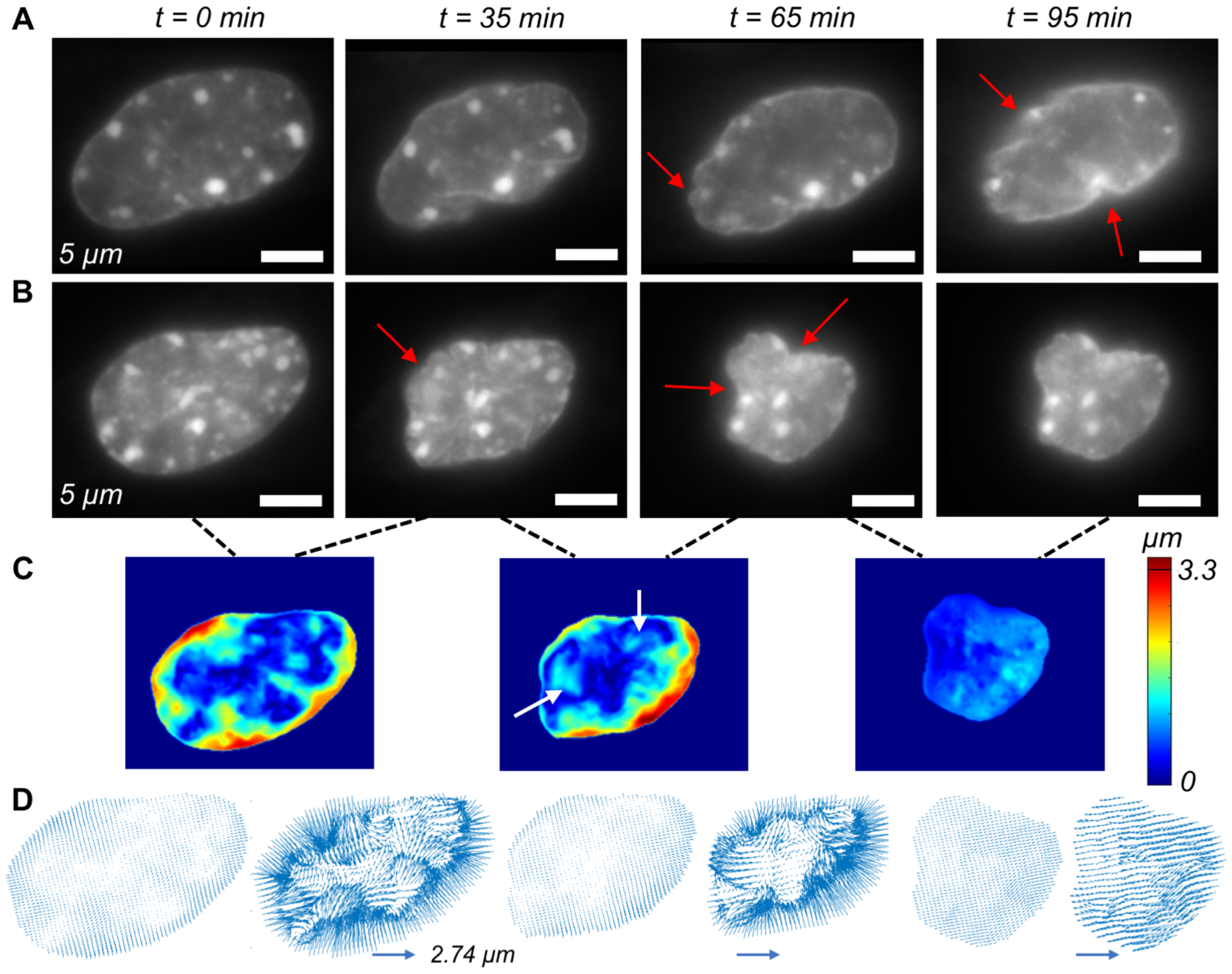
Single cell level tracking of individual Nucblue stained nuclei in the GSK126 group of NIH 3T3 cells. In all cases, the yellow arrow signifies the approximate direction of the cell migration at that timepoint. **(A)** Snapshots of a single nucleus shows that it undergoes intranuclear shape change as shown by the change in the nuclear shape and area. Nuclear ruffling, characterized by the irregular nuclear shape at the periphery is visible (red arrow). **(B)** Another example of a nucleus undergoing more extreme shape and area change. Nuclear ruffling (irregularities) is visible in the xy plane which is the plane shown the images. Nuclear ruffling is very extreme in the image corresponding to 65 min. **(C)** Rigid body motion corrected absolute displacement map of the chromatin shows the chromatin remodeling at different pairs of timesteps i.e., 0 to 35 min, 35 min to 65 min, and 65 min to 95 min. For 65 min image, rapid inflow at certain regions inside the nucleus are shown (white arrow) which is associated with the invagination (red arrow) **(D)** Vector map of the displacement field shows significant heterogeneity in the intranuclear space especially in the invaginated regions. The vector map at the right corresponding to the scale bar. The vector map in left shows scaled plots to show the exact location of the vectors, but do not represent the actual displacement magnitude.

## Discussion

The role of the cell nucleus in cell migration has only recently been elucidated. Although it is clear that the mechanics and other physiological properties of the nucleus are important in cell migration, a complete mechanistic understanding of this complex process is currently at the rudimentary stage. In this study, we showed that chromatin remodeling occurs inside the nucleus during cell migration, and with a pharmacological perturbation in chromatin remodeling, the cells lose the ability to efficiently migrate, along with disruption in nuclear shape and integrity.

Force generation by actomyosin contraction is critical for the efficient migration of cells by chemical or physical cues. As the nucleus is a relatively rigid organelle, the role of its position and mechanical properties in cell migration is not surprising. Previous studies have shown that the presence of the nucleus is critical for maintaining cell mechanical homeostasis during migration (3). However, the internal mechanics of the nucleus at the chromatin level in this process are not well understood, and this study attempts to partially fill this knowledge gap by targeted inhibition of chromatin remodeling.

This study showed that chromatin remodeling is critical for efficient cell migration. A previous study showed that in a 3D environment, TSA pretreatment softens the nucleus by blocking chromatin compaction (14) which helps cells squeeze through narrow confinements in the extracellular matrix to repopulate the scaffold during tissue engineering applications. However, that study did not focus on the cell’s ability to follow a gradient; rather, the objective was to repopulate the matrix. In the present study with 2D planar migration, we report that blocking histone deacetylase by TSA, hence hindering chromatin compaction, affects the fidelity of cell migration, as demonstrated by lower speed and random angular orientation. These data suggest that chromatin compaction at a small scale at the single nucleosome level might be required for the nucleus to be shaped by nanoscale chromatin flow and remodeling. Further insight into this process can be obtained by the selective disruption of different types of HADC proteins and their downstream epigenetic targets.

This work also suggests that chromatin remodeling at a scale larger than the single nucleosome at the multi nucleosome level is required for efficient cell migration. Blocking EZH2 with GSK126 had two effects. First, it inhibits the PRC2 (Polycomb Repressive Complex) which is critical for H2K27 trimethylation (19) Therefore, it can affect local chromatin compaction, an effect similar to that of TSA. Second, EZH2 expression is associated with ARID1A activity (20) which is known to drive chromatin remodeling as a part of the SWI/SNF complex (16). Therefore, the inhibition of EZH2 by GSK126 can affect chromatin compaction and remodeling. Our data suggest that both small-scale (single nucleosome level) and large-scale (multi nucleosome level) chromatin remodeling are critical factors in chromatin flow and remodeling. The lack of this ability in the nucleus is probably compensated for by the abrupt change in nuclear shape, which breaks down the combined homeostasis of cells and nuclear mechanics during migration. The detailed mechanism of this process can be further understood by the precise targeting of ARID1A and other proteins in SWI/SNF CRC, independent of EZH2 intervention.

It should be noted that chromatin remodeling was not stopped by GSK126 treatment (Figure 6); rather, the nucleus took an abnormal shape during migration. In addition, the non-migrating nuclei did not show any abnormal shape. This observation suggests the possibility that nuclear invagination by GSK126 treatment is a result of the requirement of the nucleus to reposition itself under inhibited chromatin remodeling capability. This could be an effect of the altered interaction between the cell and nuclear mechanics due to the altered ability of the nucleus to change its local mechanics on demand during migration. The altered homeostasis between the cell and nuclear mechanics possibly compels the nucleus to morph into an irregular shape, and it also disrupts the cell-level mechanics as well as force generation capabilities that keep the cell focused in following a chemical gradient. Further research with detailed 3D imaging of cell and nuclear structures during the migration process, along with spatiotemporal mechanical characterization in both the cytoskeleton and nucleus, can provide more physical insight into this process. The results of the present study provide insight into the complex intertwined effects of intranuclear chromatin heterogeneity, dynamic chromatin compaction, and nuclear and cell mechanics, which are mostly unknown in the context of cell migration.

## Author contributions

S.G. conceived of the project. J.F., B.H., and S.G. conducted experiments. J.F., B.H., S.K. and S.G. contributed to data analysis. S.G. wrote the manuscript with input from all authors. S.G. secured resources and funding for this study.

## Acknowledgements

The authors are grateful to the lab of Corey P. Neu at University of Colorado Boulder for providing the NIH 3T3 cells.

## Disclosure statement

No potential conflict of interest was reported by the authors.

### Data availability

The authors confirm that the data supporting the findings of this study are available in the article and its supplementary materials.

